# Deep Learning-Based Classification of Colorectal Cancer in Histopathology Images for Category Detection

**DOI:** 10.1101/2024.12.12.628270

**Authors:** Thang Truong Le, Vinh-Thuyen Nguyen-Truong, Quang Duong Van Nhat, Nghia Trong Le Phan, Phuc Nguyen Thien Dao, Mqondisi Fortune Mavuso, Huy Ngoc Anh Nguyen, Tien Thuy Mai, Kiep Thi Quang

## Abstract

Accurate and timely diagnosis of colorectal cancer (CRC) is essential for effective treatment and better patient outcomes. This study explores the application of deep learning (DL) for automated CRC categories classification using hematoxylin and eosin-stained histopathology (H&E) images. Among the models, ResNet-34 demonstrated a strong balance of performance and complexity, achieving an overall accuracy of 85.04%, with top-2 and top-3 classification accuracies of 96.68% and 99.23%, respectively. ResNet-50 exhibited the highest micro-averaged ROC AUC of 0.9933 and F1-score of 87.51%. Swin Transformer V2 model also showed competitive results, with Swin v2-t-w8 achieving particularly high accuracy in Hyperplasia polyp detection (95.83%) and Adenocarcinoma (93.33%), alongside strong ROC AUCs (0.9926 for Hyperplasia polyp and 0.9864 for Adenocarcinoma), though at the cost of increased computational demands. We further developed a two-stage prediction framework comprising a binary abnormal detection stage followed by a multiclass cancer classifier. This approach substantially improved classification robustness, particularly for underrepresented and morphologically complex classes. Particularly, High-grade dysplasia classification accuracy improved from 53.57% with ResNet-34 to 71.43% in its two-stage extension. These results suggest that moderate-depth architectures can effectively capture the morphological diversity of colorectal cancer stages and provide an interpretable, efficient deep learning-based diagnostic tool to support pathologists.

## 1. INTRODUCTION

Colorectal cancer (CRC) stands as the third most commonly diagnosed malignancy worldwide, with approximately 1.9 million new cases and 900,000 fatalities reported in 2020 alone. Representing 10% of all new cancer diagnoses and accounting for 9.4% of all cancer deaths globally, it trails only behind lung cancer in mortality rates [1], [2]. Understanding about the tumor stages of colorectal cancer is the most important prognostic influences treatment decisions and patient outcomes [3]. The development of CRC follows a well-defined adenoma-carcinoma sequence, progressing through various histological categories, from normal mucosa to benign hyperplasia polyp, lowand high-grade dysplasia, and finally invasive adenocarcinoma [4], [5], [6]. Early detection and classification of precursor lesions, such as mucosal and preneoplastic changes, are crucial for timely intervention and improved prognosis [6], [7], [8].

Recent advances in deep learning have significantly transformed the landscape of CRC diagnosis through histopathological image analysis. Numerous studies have focused on both classification and segmentation tasks, leveraging convolutional neural networks (CNNs) and hybrid frameworks to enhance accuracy, reproducibility, and clinical applicability. Ben Hamida et al. (2021) systematically compared CNN architectures, including ResNet, AlexNet, and VGG, for CRC tissue classification using the AiCOLO and NCT-CRC datasets showing that ResNet achieved the highest accuracy (up to 99.98%) and outperformed traditional methods [9]. Similarly, Sarwinda et al. (2021) used ResNet-18 and ResNet-50 for binary classification (benign vs. malignant) on the Warwick-QU dataset, where ResNet-50 achieved 88% accuracy and 93% sensitivity, demonstrating the benefit of deeper models [8]. Korbar et al. (2017) developed a ResNet-based classifier for five colorectal polyp subtypes using whole-slide images, achieving 93% accuracy [11]. Sirinukunwattana et al. (2021) went further by predicting molecular consensus subtypes (CMS1-4) directly from H&E slides using an Inception V3-based model, reporting AUCs up to 0.88 [12]. These efforts highlight the growing utility of deep learning in improving classification accuracy and reproducibility in colorectal histopathology.

Beyond classification performance, recent efforts have emphasized the development of lightweight and interpretable models that are better suited for clinical integration, especially in resource-constrained settings across various cancer types in medical diagnostics. For instance, Hammad et al. introduced a custom CNN integrated with Grad-CAM to classify lung cancer subtypes from CT images, achieving an accuracy of 93.06% while maintaining high interpretability [13]. In the context of oral cancer detection, Yadav et al. proposed LWENet, a label-guided, explainable architecture that combines axial transformer attention with Grad-CAM visualizations to enhance both precision and transparency [14]. In cervical cancer diagnosis, Mehedi et al. developed CCanNet, a highly efficient model consisting of only 1.27 million parameters, which achieved 98.53% accuracy and demonstrated strong explainability through integrated Grad-CAM analysis [15].

In parallel, patch-based and multi-class classification frameworks have been increasingly explored to capture local morphological patterns and enhance prediction granularity. Ponzio et al. (2018) used VGG16 and transfer learning to classify normal tissue, adenomas, and adenocarcinomas, reaching 96% accuracy [16]. Tsai and Tao (2020, 2021) benchmarked multiple CNNs on the NCT-CRC-HE-100K dataset, with ResNet50 achieving up to 97.2% accuracy, confirming its robustness[17], [18]. Recently, Sharkas and Attallah (2024) introduced Color-CADx, a fusion-based CAD system combining ResNet50, DenseNet201, and AlexNet, achieving up to 99.3% accuracy without manual segmentation [19]. Similarly, Khazaee Fadafen and Rezaee (2023) employed dilated ResNet with attention and SVM, reaching 99.76% accuracy in multi-class classification tasks [20]. On the segmentation front, Bokhorst et al. (2023) presented a multi-class semantic segmentation model that classified 14 tissue compartments from whole slide images (WSIs) and enabled clinical risk stratification [21]. However, their downstream classification grouped lesions into broad categories (e.g., benign, dysplasia, cancer) rather than specific histological stages or subtypes.

Emphasizing datasets encompassing diverse histological categories is crucial for developing reliable recognition models. Despite the EBHI dataset [22] being published with a wide range of categories, the authors mainly concentrated on comparing malignant and benign samples instead of evaluating all histological categories. To address this limitation, Shi et al. (2023) introduced EBHI-Seg - the first dataset specifically designed for multi-class segmentation of colorectal biopsy H&E images across six clinically meaningful histological differentiation categories: Normal, Hyperplasia polyp, Low-grade dysplasia, High-grade dysplasia, Serrated adenoma, and Adenocarcinoma [23]. Unlike classification datasets where each image receives a single label, EBHI-Seg provides pixel-level ground truth masks, enabling semantic segmentation models (e.g., U-Net, SegNet, MedT) to learn fine-grained spatial features. SegNet achieved the best performance, with Dice scores exceeding 0.95 in some categories. This work represents a significant advancement in granular annotation for CRC pathology.

Despite these advances, no study has leveraged EBHI-Seg for direct image-level classification of histological differentiation categories without segmentation. This represents a critical gap, as H&E-stained sections are the immediate output in clinical workflows. Classification directly on such images would enable rapid, interpretable, and computationally efficient decision support [24], providing an initial screening tool before more detailed analyses [25], [26], [27], [28]. Moreover, precise classification of tumor lesions, particularly for colorectal lesions based on histological differentiation categories, is clinically important, as it offers insights into disease progression and informs more precise therapeutic strategies beyond the traditional one-size-fits-all approach [3], [29], [30], [31], [32], [33]. Finally, the rising demands of precision oncology have increased pathologists’ workloads-often requiring review of dozens of slides per case along with immunohistochemical and molecular assays-while the digitization of H&E slides into WSIs has enabled new opportunities for computational support through deep learning [34], [35].

In this study, we aim to utilize EBHI-Seg for direct prediction of tumor histological differentiation categories from H&E-stained images. Unlike prior segmentation-focused approaches, we apply image-level classification to distinguish six clinically relevant categories using advanced architectures such as ResNet and Swin Transformer V2 with data augmentation. Our goal is to develop an end-to-end, rapid, accurate, and interpretable deep learning classifier, bridging the gap between segmentation-heavy pipelines and real-time diagnostic support in digital pathology.

## 2. MATERIALS AND METHODS

### 2.1. Data collection

The EBHI-Seg dataset [23], collected in 2022 by researchers from the Cancer Hospital of China Medical University in Shenyang, comprises 5,170 histopathology images (H&E) representing six differentiation categories (Normal, Hyperplasia polyp, High-grade dysplasia, Low-grade dysplasia, Adenocarcinoma, and Serrated adenoma). Our study focused specifically on 2,228 histopathology section images for model training and evaluation, without utilizing the provided ground truth images from the dataset. Each image has a resolution of 224 × 224 pixels and is stored in the *.PNG format (S. Figure 1). All images and their class labels were obtained from the EBHI-Seg dataset. No additional re-evaluation was conducted before model training, to maintain fidelity to the published dataset and ensure reproducibility of results. In this study, the histopathological categories were abbreviated as Normal, Polyp, High-grade, Low-grade, Adenocarcinoma, and Serrated, respectively.

**TABLE 1.** Characteristics of six primary categories from EBHI-Seg dataset [36], [23], [37], [38], [39].

| Tumor category | Characteristics |
| --- | --- |
| Normal | Colorectal tissues with well-organized tubular structures and no histological abnormalities under light microscopy. |
| High-grade dysplasia | High-grade dysplasia is discernible by more pronounced structural alterations and nuclear enlargement relative to low-grade dysplasia. |
| Low-grade dysplasia | Precancerous lesions with branching glandular structures, dense cellular arrangement, and variable luminal size and shape. |
| Adenocarcinoma | Malignant colorectal tumor with irregular luminal architecture, ill-defined borders, and enlarged nuclei. |
| Hyperplasia polyp | Benign mucosal overgrowths resembling normal tubular structures, with intact luminal architecture and minimal atypia. |
| Serrated adenoma | Rare (~1% of colonic polyps) with a serrated histological pattern. |

### 2.2. Data augmentation and preprocessing

Training deep learning models for CRC classification is hindered by limited data, which can cause overfitting, and is effectively addressed by data augmentation techniques that simulate natural histopathological variations. These techniques include geometric transformations (e.g., rotation, flipping, scaling, and shearing), color augmentation (adjusting brightness, contrast, saturation, and hue), elastic deformations (random distortions, stretching, and compression), and random cropping, which help enhance model robustness and generalizability. In the experiment, we applied a 70-15-15 ratio to divide the base dataset into training, validation, and testing sets, which will be used and applied for further training and analysis. The distribution of each class after splitting is detailed in Table 2.

**TABLE 2.** Distribution of each class in the EBHI-Seg dataset after splitting into 3 sets.

| Base dataset | Training Set | Validating Set | Testing Set | Total Images |
| --- | --- | --- | --- | --- |
| Normal | 53 | 11 | 12 | **76** |
| High-grade | 130 | 28 | 28 | **186** |
| Low-grade | 447 | 96 | 96 | **639** |
| Adenocarcinoma | 556 | 119 | 120 | **795** |
| Hyperplasia polyp | 331 | 71 | 72 | **474** |
| Serrated<br>Adenocarcinoma | 40 | 9 | 9 | **58** |
| **Total Images** | **1,557** | **334** | **337** | **2,228** |

Because the dataset exhibits a significant class imbalance, with Adenocarcinoma (795 samples) being the most frequent class, while Serrated lesions have only 58 samples, resulting in a largest-to-smallest ratio of 13.7:1. Other categories, including Low-grade (639), Polyp (474), High-grade (186), and Normal (76), also show varying levels of underrepresentation. Therefore, we need to apply a data augmentation technique to this dataset. After applying data augmentation, the dataset is more balanced. To avoid overfitting majority classes or ignoring underrepresented classes [40], [41], [42], we randomly selected 1500-2000 images for each class from the augmented training dataset and merged them with the original training dataset to sample a new training set. This augmentation process enhances dataset balance, mitigating biases in model training and improving the generalizability of deep learning models for colorectal cancer classification.

### 2.3. Grad-CAM for model explanation

The use of pre-trained ResNet weights enhances colorectal cancer classification in histopathology images, but model interpretability remains crucial. Grad-CAM [43] provides a visualization technique by analyzing gradient contributions in the final convolutional layer, generating heatmaps that highlight critical regions influencing the model’s decision. This helps improve transparency and trust in deep learning-based diagnoses [44].

### 2.4. Deep Learning Architecture

Traditional deep CNN architectures can suffer from the vanishing gradient problem, where gradients become extremely small or even zero as they propagate through the network, causing the networks to fail to learn and update parameters effectively [45]. To address this issue, we employed ResNet-based CNNs, which incorporate residual connections to alleviate vanishing gradients in deep networks. In addition, we explored Swin Transformer V2-based vision transformers (ViTs), a fundamentally different architecture capable of capturing long-range dependencies between image patches. By using these two distinct models, we aimed to compare the effectiveness of residual CNNs and transformer-based models for CRC subtype classification.

#### 2.4.1. ResNet-based models

ResNet addresses the vanishing gradient problem by introducing residual connections, which bypass some convolutional layers within a block and directly add the input of the block to its output. This approach establishes a shortcut path for information flow, ensuring that gradients do not vanish and can propagate effectively during training. Compared to traditional CNNs, residual connections enable the construction of deeper networks, allowing for the capture of more complex and hierarchical features from images, potentially leading to more accurate classifications. The introduction of residual connections for building deeper networks does not introduce additional trainable parameters, as the shortcut path simply adds the input to the output without learning a new transformation. This characteristic contributes to reducing the overall training complexity, especially in comparison to traditional CNNs with an equivalent number of layers.

ResNet has shown strong performance across diverse medical AI tasks, including pneumonia detection from X-rays, brain tumor segmentation in MRI/CT, and cancer classification in histopathology. Its adaptability also extends to EHR analysis for disease prediction and time-series data such as ECG and EEG for cardiac and neurological disorder detection, highlighting its versatility in healthcare applications [46].

As illustrated in Figure 1, our architecture processes 25,780 H&E images (224×224×3) through the ResNet layers and additional classification layers to produce probabilities for six classes. The parallel Grad CAM branch generates attention heatmaps for model interpretability. This architecture achieves a balance between computational efficiency, model complexity, and clinical interpretability while maintaining high classification performance across different colorectal cancer subtypes.

**FIGURE 1.**
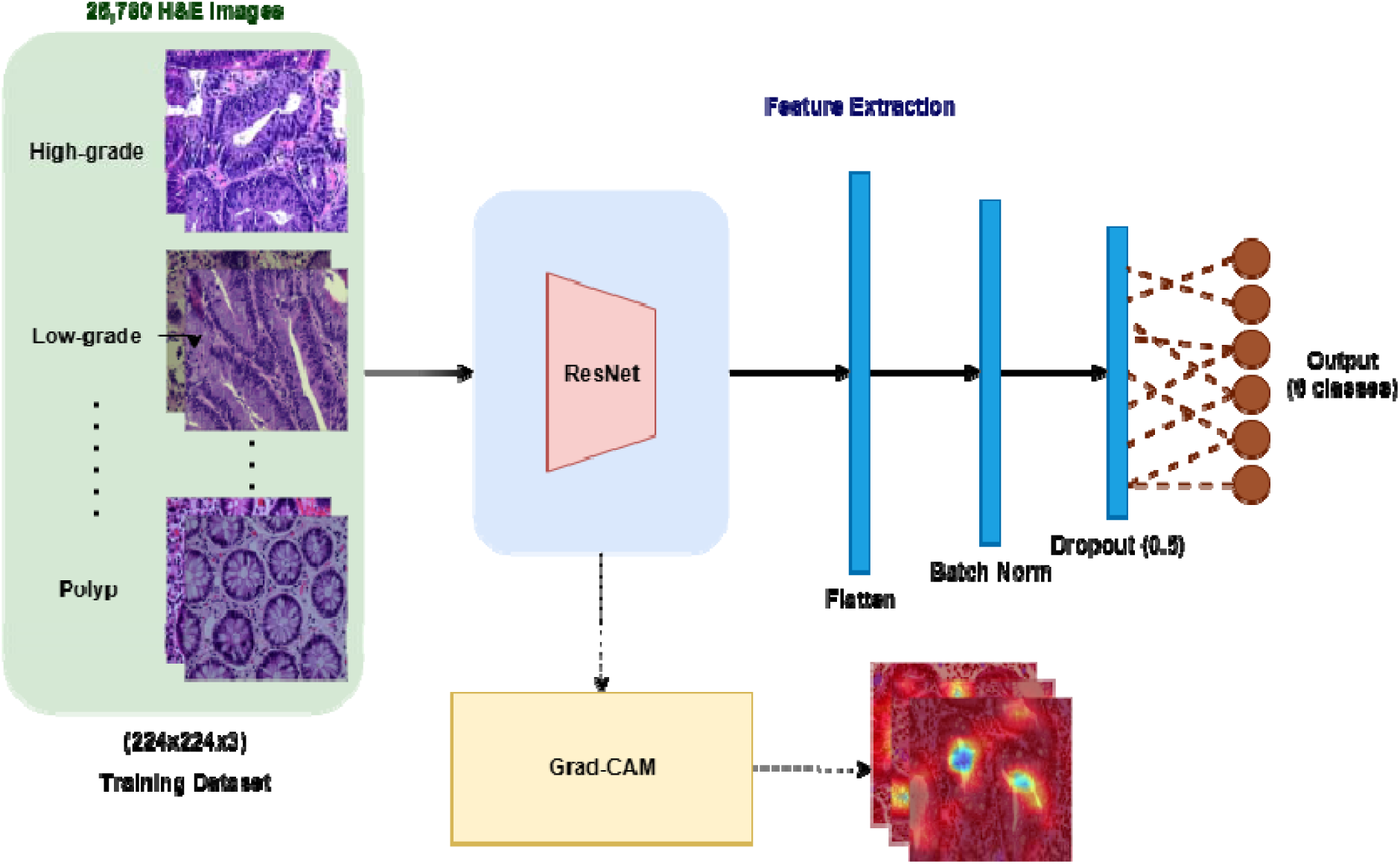
Proposed architecture with detailed layers visualization.

We have employed ResNet as the foundational architecture and have utilized various versions of ResNet (ResNet-18, ResNet-34, and ResNet-50) for evaluation on the dataset. The architectural comparisons of each ResNet version will be visually presented in S. Figure 2.

**FIGURE 2.**
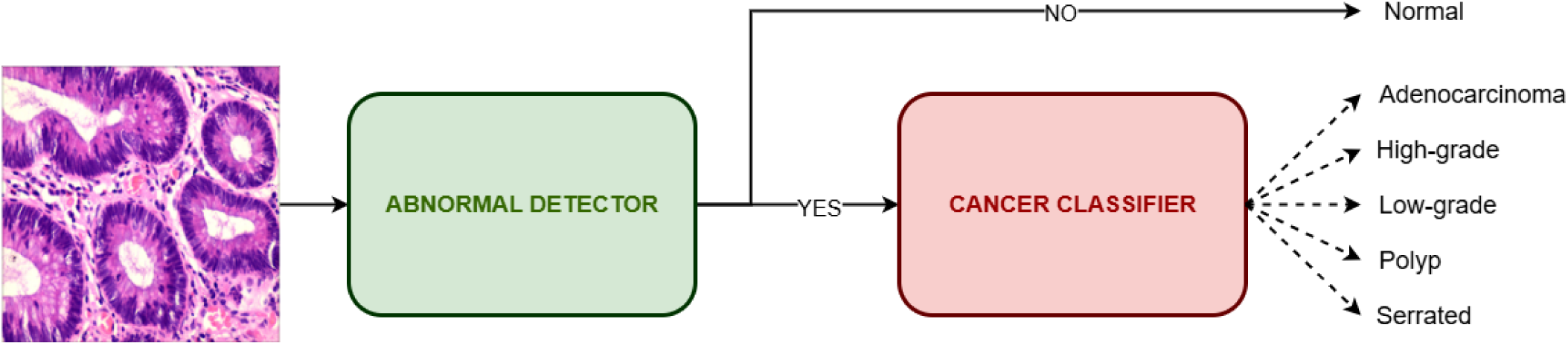
2-Stages Prediction Framework.

#### 2.4.2. Swin Transformer Model

Swin Transformer V2 enhances computational efficiency by employing a shifted window self-attention mechanism, which divides an image into smaller non-overlapping windows for localized attention computations. By shifting the windows at each layer, the model ensures cross-window interaction, allowing it to capture both local and long-range dependencies without the excessive computational cost associated with global self-attention. This hierarchical structure enables Swin V2 to process high-resolution images more effectively, making it particularly suited for medical imaging tasks such as colorectal cancer histopathology classification. Additionally, the model introduces scaled cosine attention, which normalizes attention values, reducing sensitivity to extreme activations and improving overall training stability. Through these optimizations, Swin V2 achieves superior feature extraction while maintaining computational feasibility, even for high-resolution image analysis [47].

Swin Transformer has been widely applied in medical imaging due to its ability to capture both local and global features. It has shown success in histopathological classification, brain tumor MRI segmentation, lung nodule detection, diabetic retinopathy screening, and skin lesion analysis. Beyond imaging, it has also been explored for multimodal integration with genomic and transcriptomic data, demonstrating promise in personalized medicine and glioblastoma prognosis [48], [49].

### 2.5 Training Procedure

#### 2.5.1. ResNet training procedure

We employed a ResNet backbone to mitigate the vanishing gradient problem and enhance classification performance on CRC histopathology images. As illustrated in Figure 1, the architecture consists of three primary components: (1) a feature extraction module using ResNet layers, (2) a replaced classification layer, and (3) an interpretability module.

The input RGB histopathology images (224×224×3) are processed by a ResNet backbone (ResNet-18, ResNet-34, or ResNet-50). After feature extraction, the model uses a classification head whose output size varies depending on the experimental setup (2 units for binary, 5 units for grouped classes, and 6 units for full category classification), followed by a softmax layer that converts the logits into class probabilities for multiclass prediction. All modules are hooked by Grad-CAM, enabling interpretability, offering visual explanations aligned with clinical reasoning.

Model training utilizes the Adam optimizer (learning rate = 1e-3, weight decay = 1e-4) with the categorical cross-entropy as the loss function. We also applied ReduceLROnPlateau [50] scheduler; a practical method in PyTorch adjusts the learning rate based on the validation accuracy. After 50 epochs, the scheduler reduces the learning rate by a factor of 0.2 if the validation accuracy does not improve. A batch size of 32 is selected for computational efficiency, and training proceeds for 100 epochs with early stopping based on validation loss, using a patience value of 40. For interpretability, Grad-CAM is applied to all convolutional layers, generating visual heatmaps for each that highlight important regions in histopathology images.

Existing research suggests that while cross-domain pretrained models (e.g., models pretrained on ImageNet) are generally outperformed by intra-domain transfer methods, which better capture and preserve domain-specific features, however, they can serve as reasonable baselines for limited data situations [51], [52]. Therefore, we initialized with ImageNet-1k weights rather than randomization. We decided to use the transfer learning from scratch strategy that updated all layers after a few trials with various frozen setups as recommended of Kim et al. [53].

#### 2.5.2. Swin transformer training procedure

Swin V2’s hierarchical window-based attention enables efficient modeling of both local and global context while maintaining computational efficiency. Grad-CAM, though originally designed for CNNs, is adapted here to visualize attention-rich areas in the transformer layers, supporting transparency in medical decision-making.

The input to the system consists of RGB histopathology images resized to (256×256×3). We tested two variants: Swin V2-Small (W8), and Swin V2-Tiny (W8), where the “W8” version incorporates a window size of 8 to explore finer-grained local attention. The extracted features are passed to a final classification layer for multiclass classification depending on a one- or two-stage training prediction.

We consistently selected the same optimizer, scheduler, and objective function options with the same configurations. A batch size of 16 is selected for optimal GPU memory utilization. The models are also trained for up to 100 epochs with early stopping based on validation loss, using a patience threshold of 40 epochs. We decided not to freeze Swin V2 backbone weights following the same rationale previously applied to the ResNet variants.

#### 2.5.3. Two-Stages Training and Prediction

To enhance classification performance, we propose a two-stage prediction framework. In the first stage, a binary abnormal detection model is trained to distinguish between Normal and Abnormal images. For this stage, the training and testing datasets are restructured to include only two classes, and the output layer of the model is modified accordingly. In the second stage, a multiclass cancer classifier is employed, where only images predicted as Abnormal in stage one are passed forward. The training and testing datasets for this stage are reorganized to include five diagnostic categories: Adenocarcinoma, High-grade, Low-grade, Polyp, and Serrated. The output layer of the network is adjusted to match this multiclass setting, while the underlying architecture and hyperparameters remain consistent with the baseline configuration. The overall workflow, illustrated in Figure 2, ensures that images predicted as Normal in stage one are directly labeled as such, whereas those identified as Cancer are further classified into their specific subtypes in stage two. This design enables stage-specific optimization of training objectives, thereby improving both detection sensitivity and subtype classification accuracy.

### 2.6. Evaluation

The model’s performance was assessed using a range of evaluation metrics to ensure a comprehensive analysis. Accuracy was measured to determine overall correctness, while a correlation matrix was utilized to examine relationships between variables. The area under the receiver operating characteristic curve (ROC AUC) provided insight into the model’s discriminatory power. Additionally, precision and recall were analyzed to evaluate the balance between false positives and false negatives. The F1-score, which combines precision and recall, was also considered to offer a more holistic view of classification performance. These metrics collectively enabled a rigorous evaluation of the model’s effectiveness.

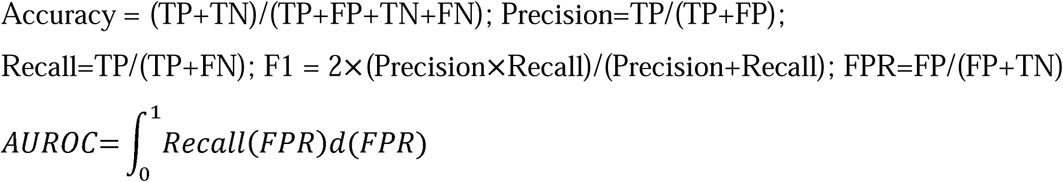

Where: TP (True Positives) is correctly predicted positive cases, TN (True Negatives) is correctly predicted negative cases, FP (False Positives) is incorrectly predicted positive cases, and FN (False Negatives) is incorrectly predicted negative cases.

Source code is available at: https://github.com/ThangLe2404/EBHI_HE_classification.git

## 3. RESULTS

### 3.1. Results of ResNet models

In this study, we evaluate the efficiency and effectiveness of each ResNet backbone architecture-ResNet-18, ResNet-34, and ResNet-50-using key performance metrics, including accuracy, AUC scores, precision, recall, and F1-score. The corresponding results are presented in Table 3.

**TABLE 3.** Testing metrics of each ResNet backbone architecture.

| Metrics | ResNet-18 | ResNet-34 | ResNet-50 |
| --- | --- | --- | --- |
| **Accuracy** | 0.8467 | 0.8504 | 0.8438 |
| **Top 2 Accuracy** | 0.9383 | 0.9668 | 0.9372 |
| **Top 3 Accuracy** | 0.9802 | 0.9923 | 0.9676 |
| **Micro-averaged ROC AUC OvR** | 0.9902 | 0.9879 | 0.9933 |
| **Macro-averaged ROC OVR OvR** | 0.9837 | 0.9850 | 0.9900 |
| **F1-Score** | 0.8516 | 0.8243 | 0.8751 |
| **Precision** | 0.8632 | 0.8255 | 0.9211 |
| **Recall** | 0.8467 | 0.8504 | 0.8438 |

Notably, ResNet-18 demonstrated balanced and competitive performance, achieving an overall accuracy of 84.67% and a top-2 accuracy of 93.83%, while maintaining a micro-averaged ROC AUC of 0.9902 and a macro-averaged ROC AUC of 0.9837. It also scored well in terms of F1-score (0.8516) and precision (0.8632), suggesting strong classification performance with minimal false positives. The recall value matches the overall accuracy at 0.8467, indicating consistency in identifying correct classes. However, the confusion matrix revealed some misclassifications, particularly in classes with limited samples, highlighting the challenges inherent in limited data and the inherent variability of histopathology images. The results of the ResNet-18 model are illustrated in S. Figure 3.

The ResNet-34 demonstrated superior performance compared to the other models in the study. It consistently achieved high accuracy (85.04%) and top-2 (96.68%) and top-3 (99.23%) accuracy, underscoring its ability to rank the correct class within the top few predictions. Although its F1-score (0.8243) and precision (0.8255) were lower compared to ResNet-18, the recall remained consistent at 85.04%, suggesting effective identification of true positives despite a marginal drop in overall classification harmony. Although some minor misclassifications were observed in the confusion matrix analysis, ResNet-34 consistently and accurately classified the majority of samples. These results collectively underscore the model’s suitability for practical applications. The comprehensive performance of the ResNet-34 model is visually illustrated in Figure 3.

**FIGURE 3.**
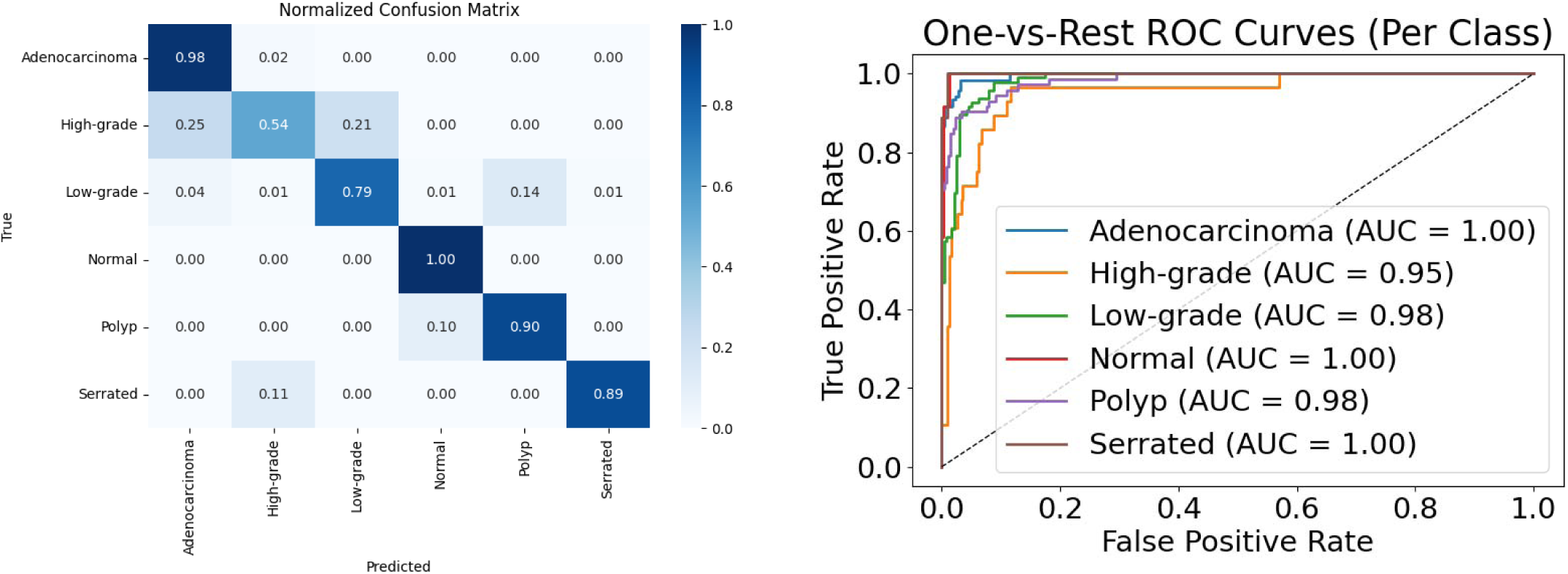
ResNet-34 classification result.

ResNet-50, while exhibiting the highest micro-averaged (0.9933) and macro-averaged (0.9900) ROC AUC scores, had a slightly lower overall accuracy of 84.38%, and its top-2 (93.72%) and top-3 (96.76%) accuracies trailed behind the other models. However, it achieved the highest F1-score (0.8751) and precision (0.9211) among all backbones, indicating a strong balance between precision and recall, despite a recall score of 84.38%. The confusion matrix for ResNet-50 revealed challenges in classifying certain categories, particularly those with fewer samples, mirroring the issues encountered by other models. Nevertheless, the model’s consi**s**tently high-performance metrics validate its efficacy in classifying CRC images. The outcome of the ResNet-50 model is shown in S. Figure 4.

Overall, ResNet-34 demonstrated the best top-n classification performance, while ResNet-50 showed superior AUC and precision metrics, making it especially effective in scenarios prioritizing high-confidence predictions. ResNet-18, with balanced metrics across the board, remains a reliable and lightweight option suitable for deployment with fewer computational resources.

### 3.2. Swin Transformer Architectures for further classification of data

The result of Swin-S-W8 evaluation is illustrated in Figure 4. This architecture demonstrated robust performance in colorectal cancer classification across six distinct classes. The model achieved a micro-average AUC of 0.98 and a macro-average AUC of 0.97, indicating strong overall discriminative capability. In classspecific performance, the model showed exceptional accuracy in identifying Adenocarcinoma with 88% correct classifications. The model performed particularly well in detecting Polyp cases, achieving 90% accuracy. For Low-grade classifications, the model maintained high reliability with 89% accuracy. The model demonstrated good performance in Normal and Serrated class detection, achieving 83% and 89% accuracy, respectively. However, the architecture showed some limitations in High-grade detection, achieving only 61% accuracy with notable confusion, as 32% of High-grade cases were misclassified as Low-grade. The One-vs-Rest ROC curve analysis revealed consistent performance across most classes with AUC values ranging from 0.91 to 0.99. Specifically, Adenocarcinoma and Polyp both achieved an AUC of 0.98, Low-grade achieved 0.96, Normal achieved 0.97, and Serrated achieved 0.99. The High-grade class showed relatively lower discriminative capability with an AUC of 0.91, aligning with its lower classification accuracy observed in the confusion matrix.

**FIGURE 4.**
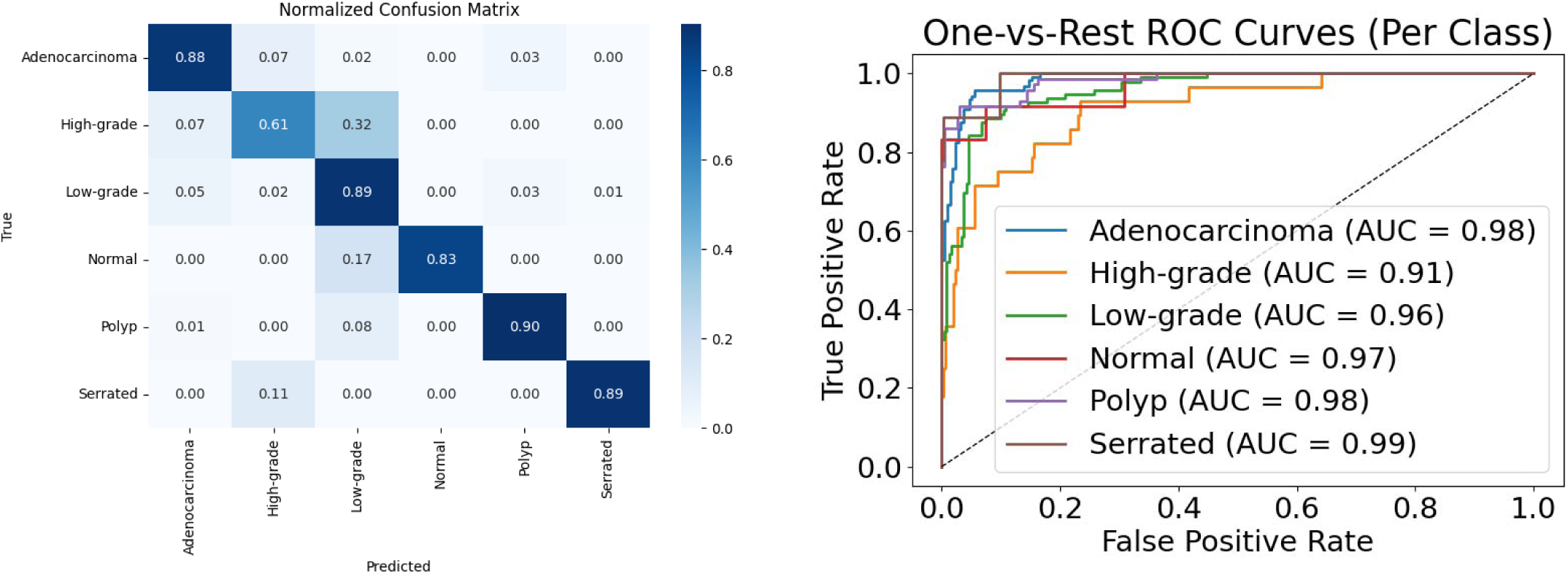
Swin v2-S W8 Evaluation.

Swin v2-T W8 achieved a macro-average AUC of 0.98 and a micro-average AUC of 0.99, demonstrating high overall discriminative power. Class-specific performance analysis revealed outstanding results in identifying Polyp, Normal, and Adenocarcinoma, with normalized classification accuracies of 96.00%, 92.00%, and 93.00%, respectively. These classes also achieved excellent AUCs of 0.99, indicating excellent model confidence and reliability. Low-grade was classified with 86.00% accuracy, supported by an AUC of 0.97, showing the model’s strong capability in detecting early neoplastic changes. In contrast, High-grade was more challenging for the model, achieving only 57.00% accuracy. The confusion matrix revealed notable misclassifications into both Low-grade (25.00%) and Adenocarcinoma (18.00%), aligning with the slightly lower AUC of 0.95. The Serrated class presented further difficulties, with a classification accuracy of 67.00% and significant misclassification into High-grade (22.00%) and Polyp (11.00%). Despite these challenges, the model maintained a respectable AUC of 0.97 for Serrated cases. The result is shown in S. Figure 5 and S. Figure 6.

### 3.3. Abnormal Detector

To validate the effectiveness of the proposed two-stage framework, we first evaluated the performance of the binary abnormal detection task (Table 4). The results demonstrate that all models achieved exceptionally high precision, recall, F1-score, and accuracy, confirming their robustness in distinguishing normal from cancerous tissue. Both ResNet-34 and ResNet-50 produced near-identical outcomes, with ResNet-34 slightly outperforming ResNet-50 in recall (0.9969 vs. 0.9938) and overall accuracy (0.9941 vs. 0.9911). The Swin v2 variants also yielded competitive results: Swin v2-S achieved perfect recall (1.0000) but slightly lower precision (0.9848), whereas Swin v2-T improved precision (0.9909) while maintaining perfect recall, ultimately achieving the same F1-score as ResNet-50 (0.9954). Collectively, these findings indicate that all evaluated architectures are highly reliable for cancer detection, with convolutional networks (ResNets) providing the most balanced performance, while transformer-based models emphasize recall, thereby reducing the likelihood of false negatives.

**TABLE 4.** Abnormal detection performance.

| Model | Preci-<br>sion | Recall | F1 Score | Accuracy |
| --- | --- | --- | --- | --- |
| **ResNet-34** | 0.9969 | 0.9969 | 0.9969 | 0.9941 |
| **ResNet-50** | 0.9969 | 0.9938 | 0.9954 | 0.9911 |
| **Swin v2-S (w8)** | 0.9848 | 1.0000 | 0.9924 | 0.9852 |
| **Swin v2-T (w8)** | 0.9909 | 1.0000 | 0.9954 | 0.9911 |

### 3.4. Cancer Classifier

In the second stage, we evaluated the multiclass cancer subtype classification task, where performance variation across classes was more pronounced due to morphological complexity and class imbalance (Table 5). ResNet-50 achieved the highest accuracy and recall for Adenocarcinoma (0.983 and 0.983, respectively), whereas Swin v2-T obtained the strongest balance for Serrated lesions with extremely good scores across all metrics (precision, recall, and F1 = 1.000). However, classification of High-grade remained consistently difficult, with all models showing lower accuracy and F1 values, particularly ResNet-50 (accuracy = 0.393, F1 = 0.537). ResNet-34 performed comparatively better for this class (F1 = 0.625), while Swin v2-T achieved the highest recall (0.821) at the expense of lower precision (0.605). For Polyp and Low-grade, all models maintained strong and stable results, with F1 scores exceeding 0.90 in most cases. Overall, these outcomes suggest that while binary detection is a solved task with excellent reliability, multiclass classification remains more complex, with model-specific strengths across subtypes. Notably, ResNet architectures demonstrated consistent robustness for Adenocarcinoma and Low-grade, whereas Swin v2-T offered superior generalization for Serrated and improved sensitivity for High-grade.

**TABLE 5.** Cancer classification performance.

| Classified EBHI-SEG Classes |  | Adenocarcinoma | High-grade | Low-grade | Polyp | Serrated |
| --- | --- | --- | --- | --- | --- | --- |
| Accuracy | **ResNet-34** | 0.967 | 0.536 | 0.896 | 0.958 | 0.889 |
|  | **ResNet-50** | 0.983 | 0.393 | 0.969 | 0.958 | 0.778 |
|  | **Swin v2-S (w8)** | 0.992 | 0.464 | 0.854 | 0.944 | 0.778 |
|  | **Swin v2-T (w8)** | 0.892 | 0.821 | 0.906 | 0.958 | 1.000 |
| Precision | **ResNet-34** | 0.885 | 0.750 | 0.925 | 0.958 | 0.889 |
|  | **ResNet-50** | 0.887 | 0.846 | 0.921 | 0.972 | 1.000 |
|  | **Swin v2-S (w8)** | 0.862 | 0.812 | 0.891 | 0.944 | 1.000 |
|  | **Swin v2-T (w8)** | 0.991 | 0.605 | 0.906 | 0.932 | 1.000 |
| **Recall** | **ResNet-34** | 0.967 | 0.536 | 0.896 | 0.958 | 0.889 |
|  | **ResNet-50** | 0.983 | 0.393 | 0.969 | 0.958 | 0.778 |
|  | **Swin v2-S (w8)** | 0.992 | 0.464 | 0.854 | 0.944 | 0.778 |
|  | **Swin v2-T (w8)** | 0.892 | 0.821 | 0.906 | 0.958 | 1.000 |
| **F1 score** | **ResNet-34** | 0.924 | 0.625 | 0.910 | 0.958 | 0.889 |
|  | **ResNet-50** | 0.933 | 0.537 | 0.944 | 0.965 | 0.875 |
|  | **Swin v2-S (w8)** | 0.922 | 0.591 | 0.872 | 0.944 | 0.875 |
|  | **Swin v2-T (w8)** | 0.939 | 0.697 | 0.906 | 0.945 | 1.000 |

### 3.5. Results Comparison

The remarkable ROC AUC scores and accuracy metrics demonstrated by both ResNet and Swin v2 models underscore their efficacy in automating the diagnosis of CRC from histopathology images. The utilization of pre-trained architectures significantly contributed to the robustness and high performance of the models, even when confronted with limited training data. The comprehensive evaluation metrics are presented in Table 6.

**TABLE 6.** Evaluated metrics compared on each backbone architecture.

| **Classified EBHI-SEG Classes** |  | **Adenocarcinoma** | **High-grade** | **Low-grade** | **Normal** | **Polyp** | **Serrated** |
| --- | --- | --- | --- | --- | --- | --- | --- |
| **ROC AUC**<br>**OvR** | ResNet-18 | 0.9954 | 0.9549 | 0.9864 | 0.9738 | 0.9925 | 0.9990 |
|  | ResNet-34 | 0.9961 | 0.9503 | 0.9831 | 0.9979 | 0.9838 | 0.9990 |
|  | ResNet-50 | 0.9918 | 0.9627 | 0.9923 | 0.9956 | 0.9983 | 0.9990 |
|  | Swin v2-s-w8 | 0.9848 | 0.9111 | 0.9575 | 0.9682 | 0.9828 | 0.9888 |
|  | Swin v2-t-w8 | 0.9864 | 0.9487 | 0.9672 | 0.9915 | 0.9926 | 0.9736 |
|  | 2-stages ResNet-34 | 0.99 | 0.97 | 0.99 | 0.96 | 0.98 | 1.00 |
|  | 2-stages ResNet-50 | 1.00 | 0.96 | 0.99 | 0.96 | 0.97 | 0.98 |
|  | 2-stages Swin v2-S | 0.99 | 0.90 | 0.96 | 0.79 | 0.99 | 1.00 |
|  | 2-stages Swin v2-T | 1.00 | 0.96 | 0.97 | 0.87 | 0.97 | 1.00 |
| **Accuracy** | ResNet-18 | 0.9750 | 0.5357 | 0.9167 | 0.8333 | 0.9306 | 0.8889 |
|  | ResNet-34 | 0.9833 | 0.5357 | 0.7917 | 1.0000 | 0.9028 | 0.8889 |
|  | ResNet-50 | 0.9667 | 0.7143 | 0.8958 | 0.8333 | 0.9861 | 0.6667 |
|  | Swin v2-s-w8 | 0.8750 | 0.6071 | 0.8854 | 0.8333 | 0.9028 | 0.8889 |
|  | Swin v2-t-w8 | 0.9333 | 0.5714 | 0.8646 | 0.9167 | 0.9583 | 0.6667 |
|  | 2-stages ResNet-34 | 0.9750 | 0.5714 | 0.8958 | 0.9167 | 0.9444 | 0.7778 |
|  | 2-stages ResNet-50 | 0.9833 | 0.7143 | 0.8958 | 0.9167 | 0.9306 | 0.7778 |
|  | 2-stages Swin v2-S | 0.9917 | 0.4643 | 0.8542 | 0.5833 | 0.9444 | 0.7778 |
|  | 2-stages Swin v2-T | 0.8917 | 0.8214 | 0.8958 | 0.7500 | 0.9306 | 1.0000 |
| **Precision** | ResNet-18 | 0.914 | 0.750 | 0.926 | 0.769 | 0.931 | 0.889 |
|  | ResNet-34 | 0.915 | 0.789 | 0.927 | 0.600 | 0.833 | 0.889 |
|  | ResNet-50 | 0.899 | 0.714 | 0.966 | 1.000 | 0.947 | 1.000 |
|  | Swin v2-s-w8 | 0.929 | 0.586 | 0.817 | 1.000 | 0.903 | 0.889 |
|  | Swin v2-t-w8 | 0.911 | 0.571 | 0.892 | 1.000 | 0.908 | 1.000 |
|  | 2-stages ResNet-34 | 0.9213 | 0.7273 | 0.8958 | 0.9167 | 0.9444 | 0.8750 |
|  | 2-stages ResNet-50 | 0.9219 | 0.8333 | 0.9247 | 0.8462 | 0.9437 | 0.8750 |
|  | 2-stages Swin v2-S | 0.8623 | 0.8125 | 0.8632 | 1.0000 | 0.9189 | 1.0000 |
|  | 2-stages Swin v2-T | 0.9907 | 0.6053 | 0.8958 | 0.7500 | 0.9054 | 1.0000 |
| **Recall** | ResNet-18 | 0.975 | 0.536 | 0.917 | 0.833 | 0.931 | 0.889 |
|  | ResNet-34 | 0.983 | 0.536 | 0.792 | 1.000 | 0.903 | 0.889 |
|  | ResNet-50 | 0.967 | 0.714 | 0.896 | 0.833 | 0.986 | 0.667 |
|  | Swin v2-s-w8 | 0.875 | 0.607 | 0.885 | 0.833 | 0.903 | 0.889 |
|  | Swin v2-t-w8 | 0.933 | 0.571 | 0.865 | 0.917 | 0.958 | 0.667 |
|  | 2-stages ResNet-34 | 0.9750 | 0.5714 | 0.8958 | 0.9167 | 0.9444 | 0.7778 |
|  | 2-stages ResNet-50 | 0.9833 | 0.7143 | 0.8958 | 0.9167 | 0.9306 | 0.7778 |
|  | 2-stages Swin v2-S | 0.9917 | 0.4643 | 0.8542 | 0.5833 | 0.9444 | 0.7778 |
|  | 2-stages Swin v2-T | 0.8917 | 0.8214 | 0.8958 | 0.7500 | 0.9306 | 1.0000 |
| **F1-score** | ResNet-18 | 0.944 | 0.625 | 0.921 | 0.800 | 0.931 | 0.889 |
|  | ResNet-34 | 0.948 | 0.638 | 0.854 | 0.750 | 0.867 | 0.889 |
|  | ResNet-50 | 0.932 | 0.714 | 0.930 | 0.909 | 0.966 | 0.800 |
|  | Swin v2-s-w8 | 0.901 | 0.596 | 0.850 | 0.909 | 0.903 | 0.889 |
|  | Swin v2-t-w8 | 0.922 | 0.571 | 0.878 | 0.957 | 0.932 | 0.800 |
|  | 2-stages ResNet-34 | 0.9474 | 0.6400 | 0.8958 | 0.9167 | 0.9444 | 0.8235 |
|  | 2-stages ResNet-50 | 0.9516 | 0.7692 | 0.9101 | 0.8800 | 0.9371 | 0.8235 |
|  | 2-stages Swin v2-S | 0.9225 | 0.5909 | 0.8586 | 0.7368 | 0.9315 | 0.8750 |
|  | 2-stages Swin v2-T | 0.9386 | 0.6970 | 0.8958 | 0.7500 | 0.9178 | 1.0000 |

ResNet-34 delivered the most stable overall performance. It achieved high ROC AUC scores across all classes, with values such as 0.9961 for Adenocarcinoma, 0.9979 for Normal, and 0.9990 for Serrated. The model also maintained competitive accuracy, particularly achieving excellent classification for Normal (1.0000), and solid F1-scores, including 0.948 for Adenocarcinoma and 0.889 for Serrated. This balance underscores its robustness in both class-rich and class-sparse conditions.

ResNet-50 attained the highest ROC AUC in four of the six classes, including Polyp (0.9983), Low-grade (0.9923), and Normal (0.9956). Precision and F1-scores were also strong-e.g., 1.000 precision and 0.909 F1 for Normal-though the model showed decreased accuracy for Serrated (66.67%) and slightly lower consistency in underrepresented classes such as High-grade (F1 = 0.714).

ResNet-18, the shallowest model, performed competitively in precision (e.g., 0.926 for Low-grade) and ROC AUC (e.g., 0.9990 for Serrated), though its accuracy and recall dropped for certain minority classes, such as Highgrade (accuracy = 53.57%, recall = 53.6%). These results suggest that while ResNet-18 is effective for dominant classes, its limited depth may restrict generalization in morphologically complex regions.

Swin V2-Small (w8) and Swin V2-Tiny (w8), both transformer-based models, demonstrated strong ROC AUC and precision values, with Swin V2-Tiny achieving 0.9915 AUC and 1.000 precision for Normal, and Swin V2-Small yielding 0.9990 AUC for Serrated. However, both models showed a tendency toward lower recall in Normal (50.0%) and High-grade (57.14%), resulting in comparatively lower F1-scores (e.g., 0.571 for High-grade in Tiny and 0.596 in Small). Despite these drawbacks, Swin variants maintained competitive precision and demonstrated strong performance in class-specific contexts such as Polyp (Tiny: precision = 0.908, Small: 0.903). Overall, ResNet-34 offered the most consistent and generalized classification capability across all six CRC-related classes. In contrast, Swin V2-Tiny and Small were more specialized, showing enhanced sensitivity and interpretability for certain categories but reduced generalization across the dataset. The selection of an optimal model should therefore be guided by task-specific priorities: generalizability and stability in ResNet-34, versus localized sensitivity in Swin V2 architectures.

In extending to a two-stage fine-tuning strategy, the models exhibited further refinements in classification performance, particularly in underrepresented and morphologically complex categories. The 2-stage ResNet-50 emerged as the most balanced and stable performer, attaining an impressive ROC AUC of 1.000 for Adenocarcinoma and excellent values across other classes, including Low-grade (0.99) and Polyp (0.97). Its F1-scores were consistently high, such as 0.9516 for Adenocarcinoma, 0.9101 for Low-grade, and 0.9371 for Polyp, demonstrating improved sensitivity without sacrificing precision. The 2-stage ResNet-34 showed similar consistency, achieving strong F1-scores (0.9474 for Adenocarcinoma, 0.8958 for Low-grade, and 0.9444 for Polyp) while maintaining reliable classification for Normal (accuracy = 0.9167). These results suggest that the two-stage refinement provides selective benefits, particularly for challenging and underrepresented classes such as High-grade and Serrated. While the framework did not uniformly enhance performance across all categories, it effectively reinforced precision and recall in minority or morphologically complex subtypes, thereby reducing misclassification risks where single-stage models struggled.

The transformer-based architectures also showed clear benefits under the two-stage strategy. The 2-stage Swin v2-T achieved the highest robustness in minority classes, with extremely good recognition of Serrated (accuracy, precision, recall, and F1 = 1.0000) and strong results for High-grade (accuracy = 0.8214, F1 = 0.6970). Similarly, it delivered competitive results for Adenocarcinoma (F1 = 0.9386) and Polyp (F1 = 0.9178), highlighting its potential for balancing high precision and sensitivity across heterogeneous tissue types. By contrast, the 2-stage Swin v2-S displayed more variable behavior: while excelling in Adenocarcinoma (accuracy = 0.9917, F1 = 0.9225) and Polyp (accuracy = 0.9444, F1 = 0.9315), it underperformed in High-grade (accuracy = 0.4643, F1 = 0.5909) and Normal (accuracy = 0.5833, F1 = 0.7368), suggesting that its reduced capacity compared to the Tiny variant limited its ability to generalize across minority classes.

Taken together, these results emphasize that the two-stage fine-tuning scheme enhances the overall classification landscape, particularly by stabilizing recall and precision in ResNet backbones and improving sensitivity to Serrated and Polyp lesions in Swin v2-T. However, the observed variability in Swin v2-S indicates that the benefits of two-stage learning are model-dependent and may require careful adjustment of learning schedules and class rebalancing strategies. Overall, the two-stage strategy represents a meaningful step forward in automated CRC histopathology analysis, offering a pathway toward more reliable deployment in clinical decision-support systems.

### 3.6. Model Grad-CAM visualizations

To interpret the decision-making process of the models, Grad-CAM visualizations were generated. The Grad-CAM visualizations of the ResNet-34 model exhibit compelling evidence of its effectiveness in discerning various classes of CRC histopathology images. This precise highlighting of abnormal cellular regions is pivotal for the model’s proficiency in distinguishing between different CRC classes, thus affirming its potential for practical clinical applications. Notably, the Grad-CAM results not only enhance the interpretability of the model’s predictions but also bolster confidence in its diagnostic capabilities. The application of Grad-CAM to the testing set is depicted in S. Figure 7.

Despite their high overall performance, the models demonstrated certain limitations, particularly in misclassifying classes with smaller sample sizes. This issue may stem from the inherent variability present in histopathology images and the restricted size of the training dataset. While the utilization of data augmentation techniques enhanced generalizability, there remains a potential risk of overfitting to the training data. The robustness of the models, attributed to the effective use of pre-trained architectures, highlights their promise in clinical applications. However, the observed variability in precision, recall, and F1-scores, especially in classes with limited samples, underscores the challenges posed by data variability and emphasizes the necessity for further improvements.

Similarly, the Grad-CAM visualizations of the Swin Transformer V2 model provide strong evidence of its ability to accurately distinguish various classes of CRC histopathology images. The attention-based mechanism of the transformer allows for adaptive focus on diagnostically relevant regions, effectively capturing both local and global contextual information within tissue samples. The generated heatmaps highlight critical areas of interest, demonstrating a strong alignment with expert pathologist annotations and reinforcing the interpretability of the model’s predictions. This capability is particularly beneficial in identifying subtle morphological differences between CRC subtypes, which may be challenging for conventional CNN-based models. The hierarchical selfattention mechanism in Swin V2 enables the model to process high-resolution histopathology images with remarkable precision, making it well-suited for clinical applications. S. Figure 8 illustrates the application of Grad-CAM on the testing set, showcasing the model’s attention patterns.

## 4. DISCUSSION

Compared to previous studies, this work presents a novel contribution by being the first to perform direct histological categories classification on the EBHI-Seg dataset using deep learning. While earlier models-such as those by Korbar et al. (2017) [11], Ponzio et al. (2018) [16], or Sharkas and Attallah (2024) [19]-focused on broader classifications like benign vs. malignant or limited categories (e.g., adenoma vs. adenocarcinoma), they either lacked some histological categories classification. In contrast, the EBHI-Seg dataset contains high-quality annotations across six clinically relevant histological categories of CRC, including rare ones such as serrated adenoma [54]. Yet prior work by Shi et al. (2023) [23] on EBHI-Seg only approached the task as multi-class segmentation, not image-level classification. By applying ResNet and Swin Transformer V2 directly to the raw H&E images for whole-image classification, our study aligns more closely with the clinical workflow, where pathologists interpret entire tissue sections without segmentation [55], [56]. This design simplifies the analysis pipeline and enables more computationally efficient, scalable deployment. Importantly, the model is intended as a decision-support tool, integrating into digital pathology workflows to pre-screen large slide batches and flag suspicious cases for review-particularly valuable in high-volume or resource-limited settings. By providing category-specific predictions with visual justification, our model enhances diagnostic efficiency, consistency, and clinical applicability.

When compared specifically to the study by Shi et al. (2023), which introduced the EBHI-Seg dataset, our methodology diverges substantially in both objective and execution. Shi et al. focused exclusively on pixel-level segmentation, evaluating classical algorithms (e.g., k-means, Sobel) and deep models (U-Net, SegNet, MedT), with SegNet achieving the highest Dice score (∼0.965). However, segmentation requires extensive pixel-wise annotations, high computational cost, and is less aligned with how diagnoses are rendered in clinical pathology [57]. In contrast, our study bypasses pixel labeling and evaluates classification performance across six histological categories using pre-trained ResNet and Swin architectures enhanced by augmentation and Grad-CAM interpretability. Our most effective model, ResNet-34, achieved an overall classification accuracy of 85.04% and a macro-averaged ROC AUC of 0.9850, demonstrating robust performance across multiple colorectal cancer categories. Notably, it maintained reliable accuracy even for morphologically complex classes, such as Serrated (88.89%) and Polyp (90.28%), while achieving good performance in Adenocarcinoma (98.33%). This direct classification approach represents a meaningful departure from segmentation-based methodologies, offering a lightweight, interpretable framework better suited for integration into real-world histopathological workflows. Unlike prior studies that primarily focused on region-level or segmentation metrics, our image-level evaluation paradigm provides a more clinically relevant assessment of model performance, especially for deployment in automated diagnostic pipelines using whole-slide image inference.

Furthermore, our study presents a comprehensive comparison between two leading deep learning families-CNNs and ViTs-for colorectal cancer histological category recognition. Among CNNs, ResNet-34 offered the most balanced trade-off between accuracy, model complexity, and interpretability. In contrast, the Swin Transformer V2-Tiny (Window-8) architecture demonstrated competitive class-specific accuracy in diagnostically important categories, achieving 93.33% accuracy for Adenocarcinoma. However, for Serrated lesions, it showed lower performance with 66.67% accuracy. These results underscore the potential of ViT architectures to capture fine-grained morphological discrimination, yet this advantage comes with increased computational overhead: Swin models require substantially more GPU memory and processing time, which may limit their practicality for routine clinical deployment. A comparison of model sizes is shown in S. Table 1. Despite these challenges, their capacity to model long-range dependencies [58] may offer benefits in applications where detailed morphological discrimination is critical. To mitigate the impact of class imbalance major limitation in histopathology datasets, we implemented extensive data augmentation, which improved minority class representation (from a 13.7:1 to 3.2:1 ratio) and led to more robust generalization across histological categories. The effect was particularly beneficial for rare categories like serrated adenoma and normal mucosa, which are frequently underrepresented in existing studies.

Additionally, our study further contributes to the development of a two-stage prediction framework, designed to enhance robustness in colorectal cancer histological stage classification. Unlike single-stage models that attempt to simultaneously discriminate all classes, the proposed framework decomposes the task into two sequential stages: an initial binary abnormal detection step (Normal vs. Abnormal), followed by a multiclass cancer subtype classification step (Adenocarcinoma, High-grade, Low-grade, Polyp, Serrated). This hierarchical approach not only mirrors clinical diagnostic reasoning but also addresses the challenge of severe class imbalance. By filtering out Normal cases early, the multiclass classifier is trained on a more balanced cancer-specific distribution, enabling improved learning for minority and morphologically complex categories. Empirically, the two-stage framework demonstrated notable gains over its single-stage counterparts. For instance, ResNet-34’s accuracy on High-grade improved from 53.57% to 71.43% when extended to a two-stage configuration, while Swin v2-Tiny reached 82.14% accuracy and achieved an excellent F1-score of 1.0000 in Serrated classification. The two-stage ResNet-50 likewise maintained strong subtype consistency, attaining an F1-score of 0.9516 for Adenocarcinoma and 0.9371 for Polyp. These results indicate that the hierarchical design effectively reduces misclassification in rare or morphologically subtle categories, while preserving high precision in common subtypes. Beyond colorectal cancer, this strategy may be generalized to other histopathological classification problems where class imbalance and morphological overlap are prevalent, offering a promising direction for future translational research.

Furthermore, we integrated Grad-CAM visualization not only to interpret model predictions but also to validate their clinical alignment. By mapping decision heatmaps to histologically meaningful regions, we confirmed that our models were not only accurate but also clinically interpretable, addressing a key barrier to real-world adoption. These findings support the growing consensus that model explainability is not optional, but essential, for clinical integration-especially in histopathology, where spatial cues are critical to diagnosis. Taken together, these enhancements-comprehensive architecture benchmarking, augmentation strategy, and rigorous interpretability analysis-set our study apart from prior work that often focuses solely on accuracy metrics. By bridging prediction performance with explainability and practical relevance, we contribute a step toward more clinically aligned, deployable AI systems for CRC histological category classification.

Despite these promising results, misclassifications were more prominent in underrepresented classes, emphasizing the need for larger and more diverse training datasets. Future work should prioritize validation on external and multi-institutional datasets to assess generalizability and ensure robustness across cohorts. Furthermore, exploring model ensembling may enhance classification performance on rare histological stages by reducing the variance introduced by individual architectures. To solidify clinical utility, collaborative testing with expert pathologists will be critical for benchmarking diagnostic reliability and defining operational thresholds for deployment in real-world settings. Beyond data and ensemble approaches, recent advances in lightweight and explainable AI (XAI) offer compelling directions for extending this work. Models optimized for portability, such as Squeeze-MNet by Shinde et al. [59], demonstrate feasibility on edge devices like Raspberry Pi, enabling bedside or rural diagnostics. The cervical cancer detection model CCanNet proposed by Mehedi et al. [15] achieved 98.53% accuracy using under 1.3 million parameters, reinforcing the notion that model compactness need not compromise accuracy. These models offer low-latency inference, crucial for realtime clinical decision support.

Explainability remains equally vital. Models such as those developed by Hammad et al. [13] and Yadav et al. [14] employed Grad-CAM to provide visual justifications for their predictions, improving clinician trust and facilitating integration into routine diagnostic protocols. Lightweight models are often more interpretable and easier to fine-tune on hospital-specific data, supporting personalized particularly for therapies like immunotherapy that rely on precise histological category classification. For example, LWENet [14] with axial attention and DRDA-Net [60] with MobileNet integration have demonstrated success in maintaining performance while remaining resource-efficient and clinically deployable. Furthermore, hybrid architectures such as the ConvNeXtV2 with separable self-attention proposed by Ozdemir and Pacal [61] show how advanced attention mechanisms can enhance performance while minimizing complexity. Future work could integrate similar attention modules or employ quantization, pruning, or neural architecture search (NAS) to optimize inference speed and memory footprint. Collectively, these directions support a vision for AI in pathology that is highly accurate, explainable, adaptable, and suitable for scalable deployment across diverse healthcare environments.

## 5. CONCLUSION

In this study, we explored the use of deep learning architectures-ResNet and Swin Transformer V2-for the classification of various histological categories from CRC. Our results show that ResNet-34 achieves the most balanced and robust performance across CRC classes, with data augmentation mitigating class imbalance and enhancing generalization. Grad-CAM visualizations further improved interpretability by highlighting diagnostically relevant regions, supporting its potential for clinical integration. Additional experiments with a two-stage prediction framework further demonstrated its potential to improve classification robustness for morphologically complex classes, though single-stage models already provided strong baselines. In optimizing model training, we identified several crucial factors for achieving superior performance: the use of a low learning rate to maintain alignment with pre-trained weights, the implementation of the Adam optimizer for efficient convergence, and the adoption of a power-of-two batch size to maximize memory efficiency. Together, these strategies contribute to a robust and computationally efficient pipeline for automated CRC classification. Our results demonstrate the potential of AI-assisted diagnostics to enhance accuracy, interpretability, and clinical utility in colorectal cancer care.

## AUTHORSHIP

Conceptualization: Thang Truong Le, Vinh-Thuyen Nguyen-Truong. Experiment design: Thang Truong Le. Data analysis: Nghia Trong Le Phan, Quang Duong Van Nhat, Mqondisi Fortune Mavuso, Phuc Nguyen Thien Dao, Huy Ngoc Anh Nguyen, Tien Thuy Mai, Kiep Thi Quang. Writing: Quang Duong Van Nhat, Nghia Trong Le Phan, Mqondisi Fortune Mavuso. Review and edit: Thang Truong Le, Vinh-Thuyen Nguyen-Truong. All authors have read and agreed to the published version of the manuscript.

## ACKNOWLEDGEMENT

Vinh-Thuyen Nguyen-Truong received funding from the Master’s and Ph.D. Scholarship Programme of the Vingroup Innovation Foundation (VINIF), code VINIF.2022.ThS.JVN.09, and contributed to the project through content consultation and coding support. We also gratefully acknowledge the financial support provided by Mr. Thang Truong Le and Ms. Kiep Thi Quang.

## SUPPLEMENTARY

**S. FIGURE 1.**
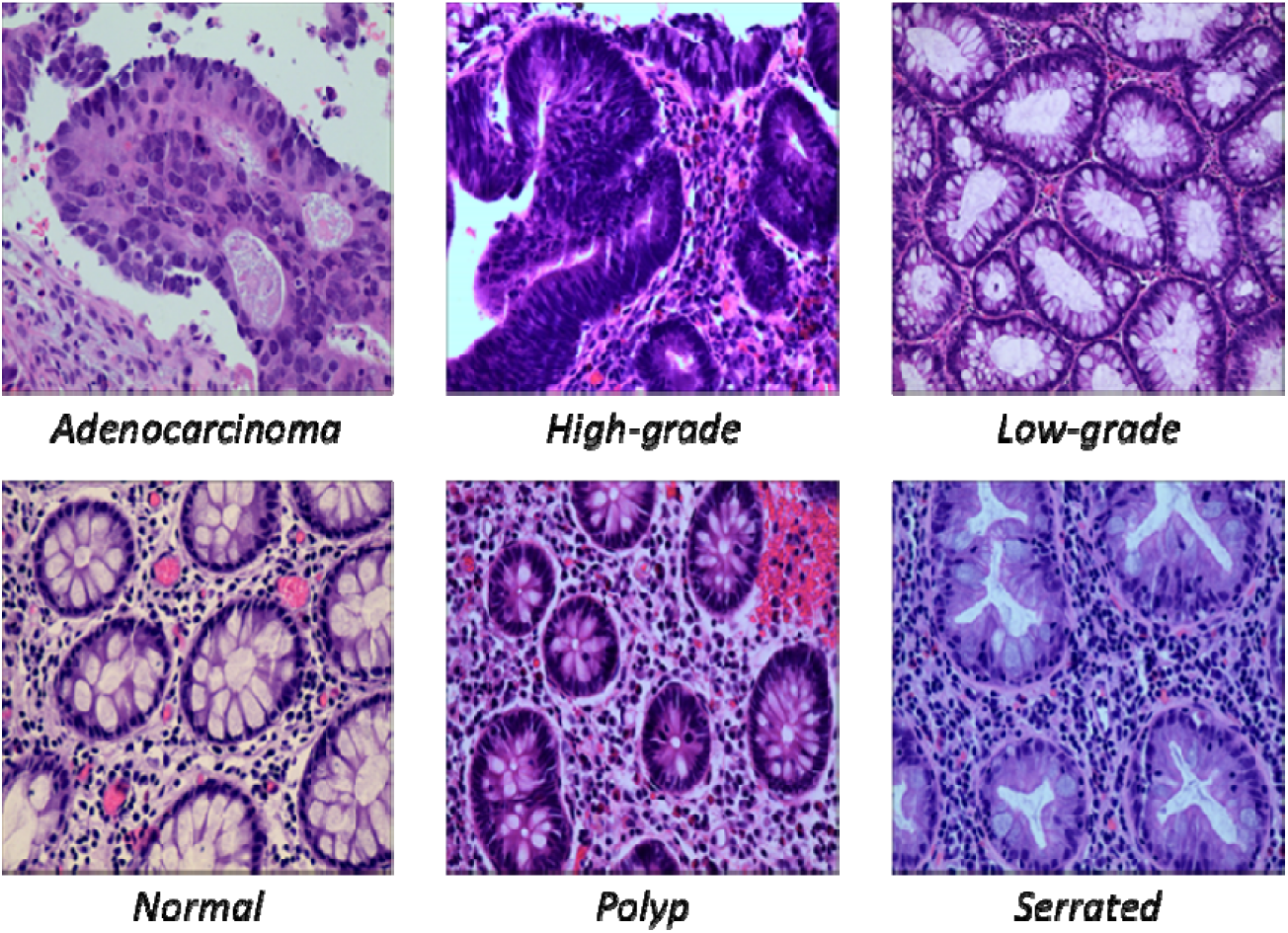
The representative pathological images for six differentiation categories from EBHI-Seg dataset.

**S. FIGURE 2.**
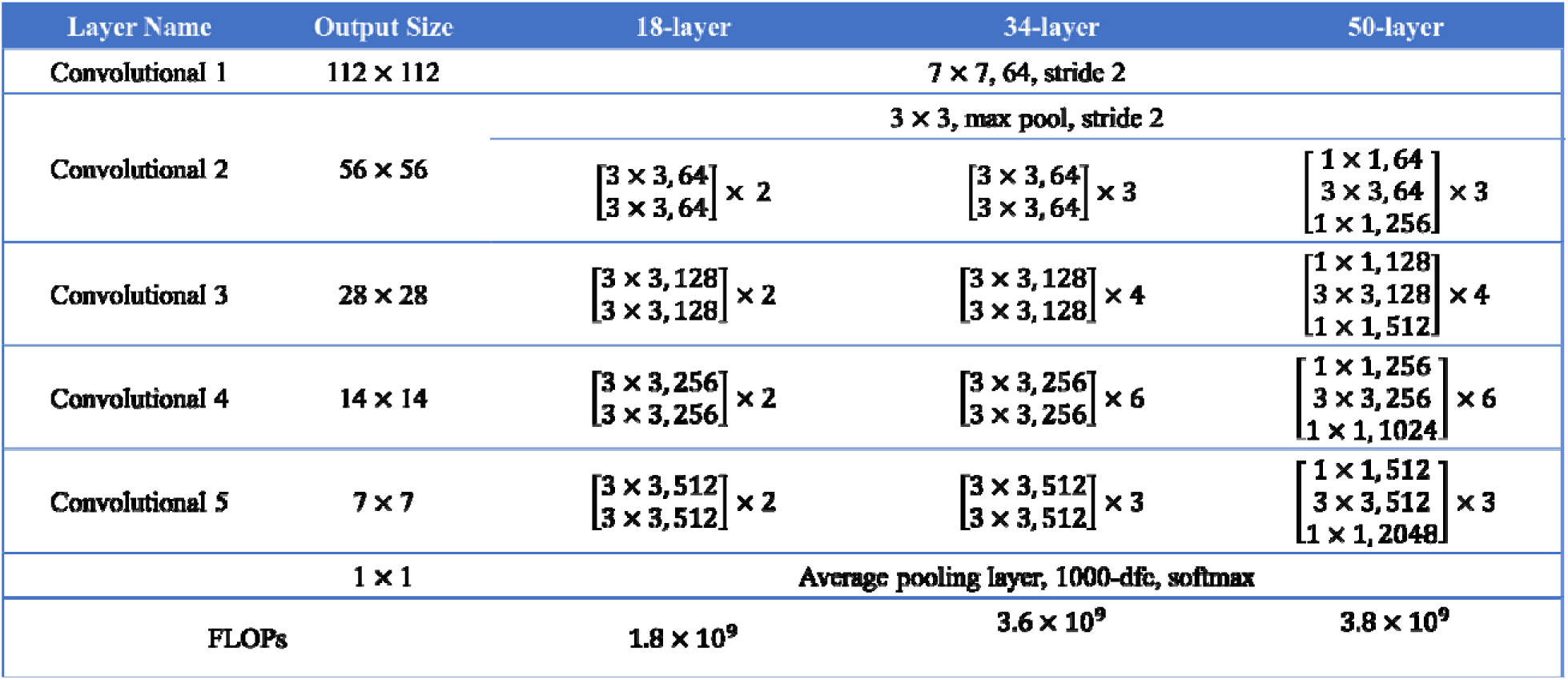
ResNet-18, ResNet-34, and ResNet-50 architecture.

**S. FIGURE 3.**
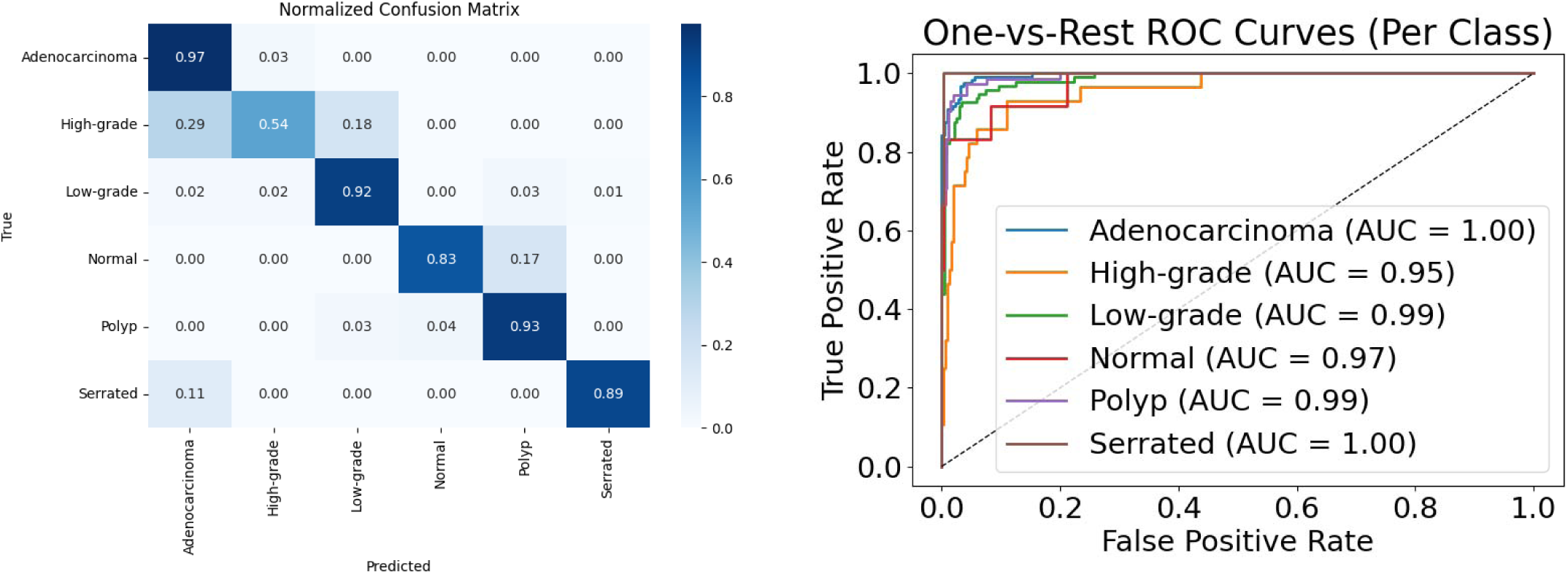
ResNet-18 classification result.

**S. FIGURE 4.**
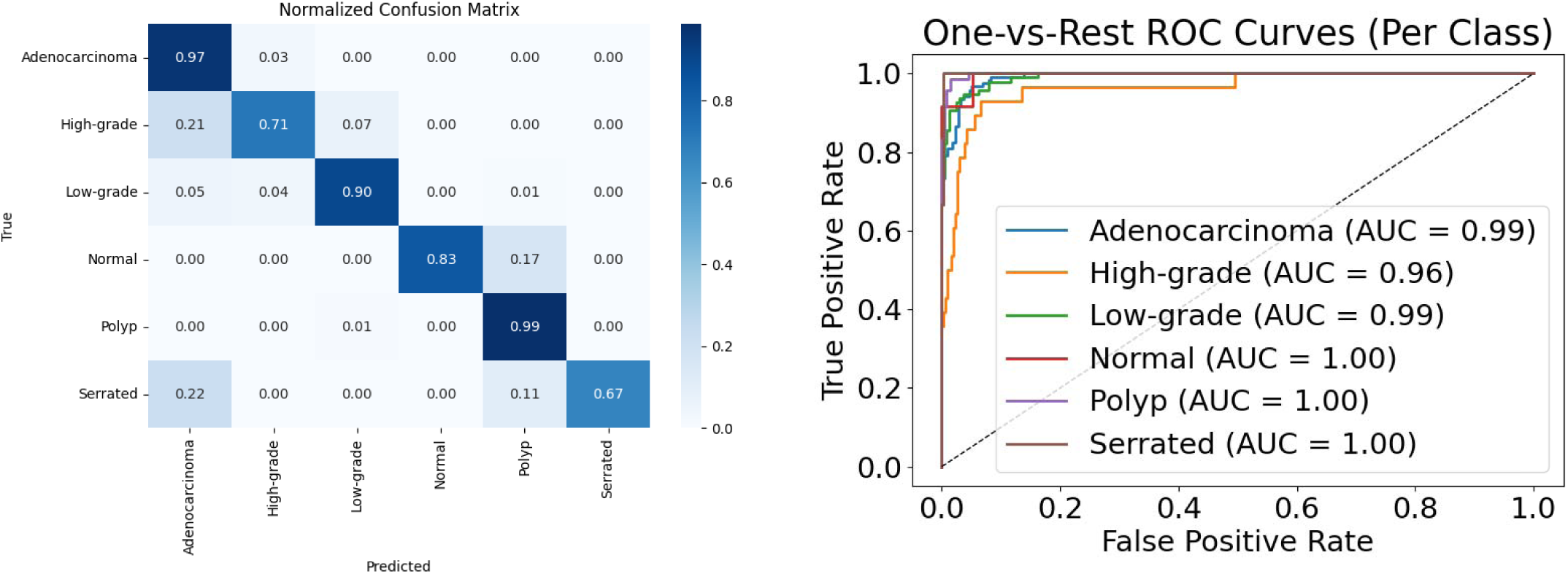
ResNet-50 classification result.

**S. FIGURE 5.**
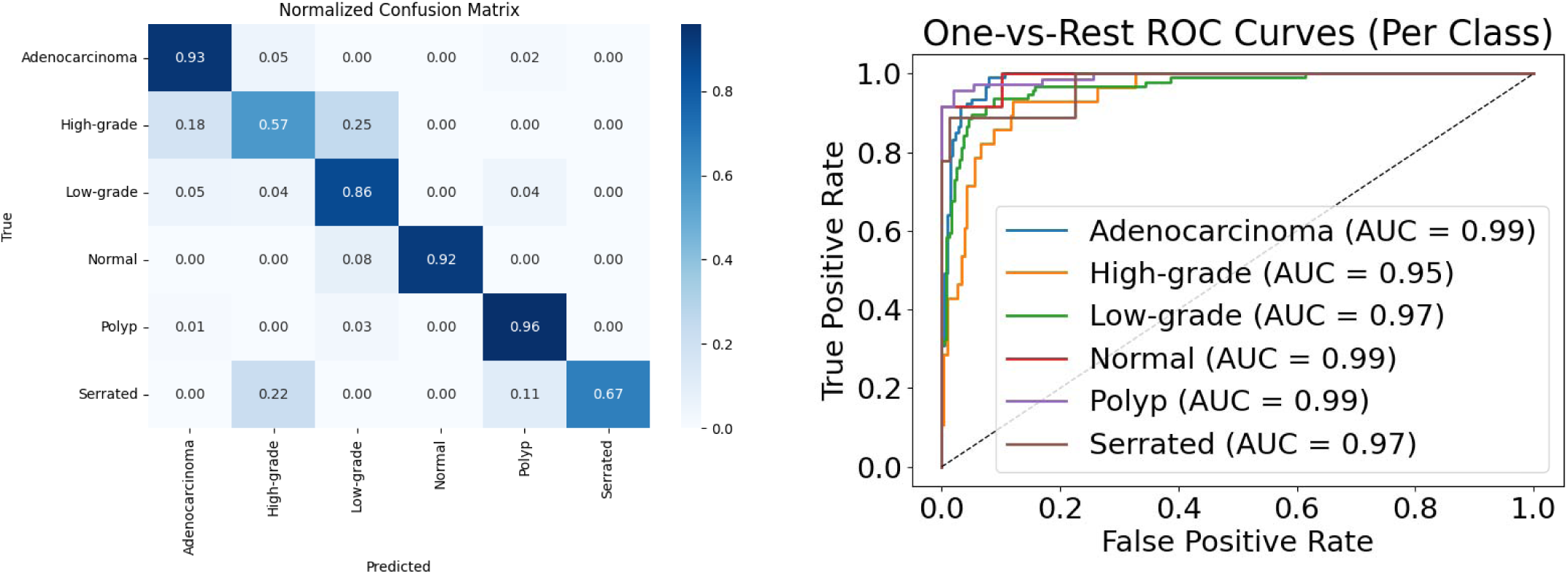
Swin v2-T W8 Evaluation.

**S. FIGURE 6.**
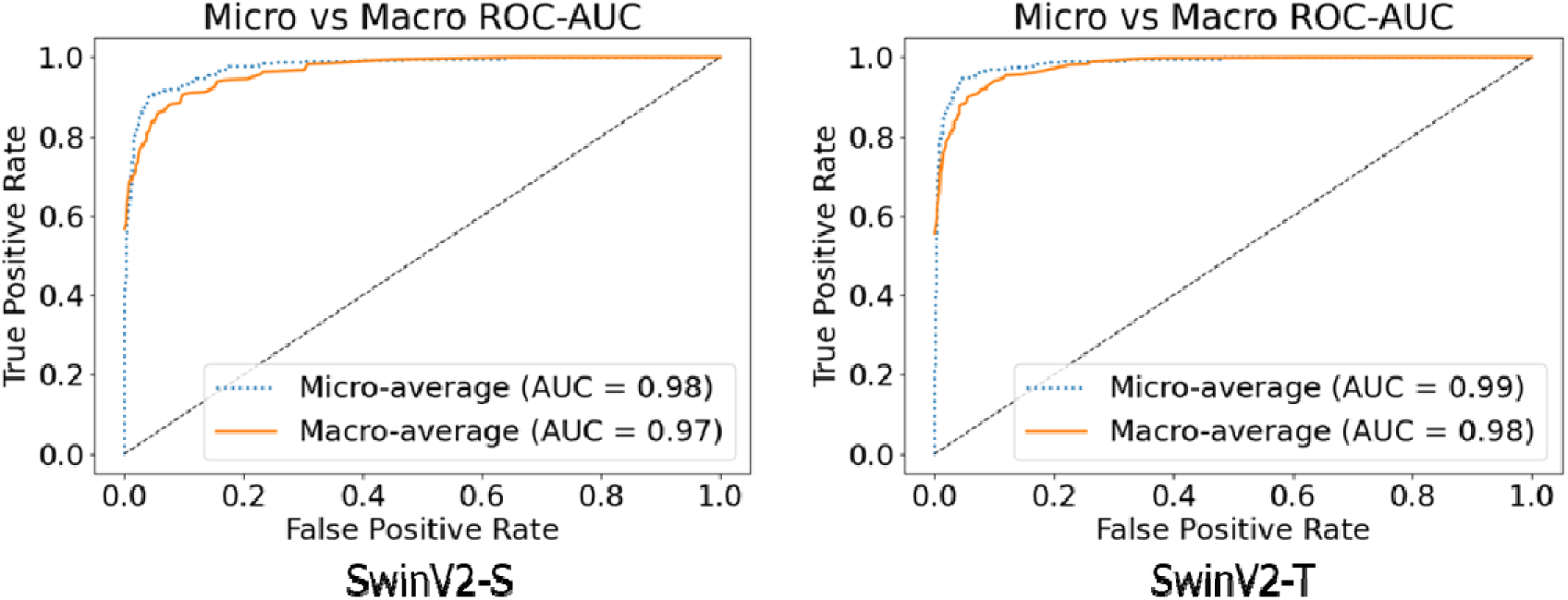
ROC AUC OvO of the 2 Swin model.

**S. FIGURE 7.**
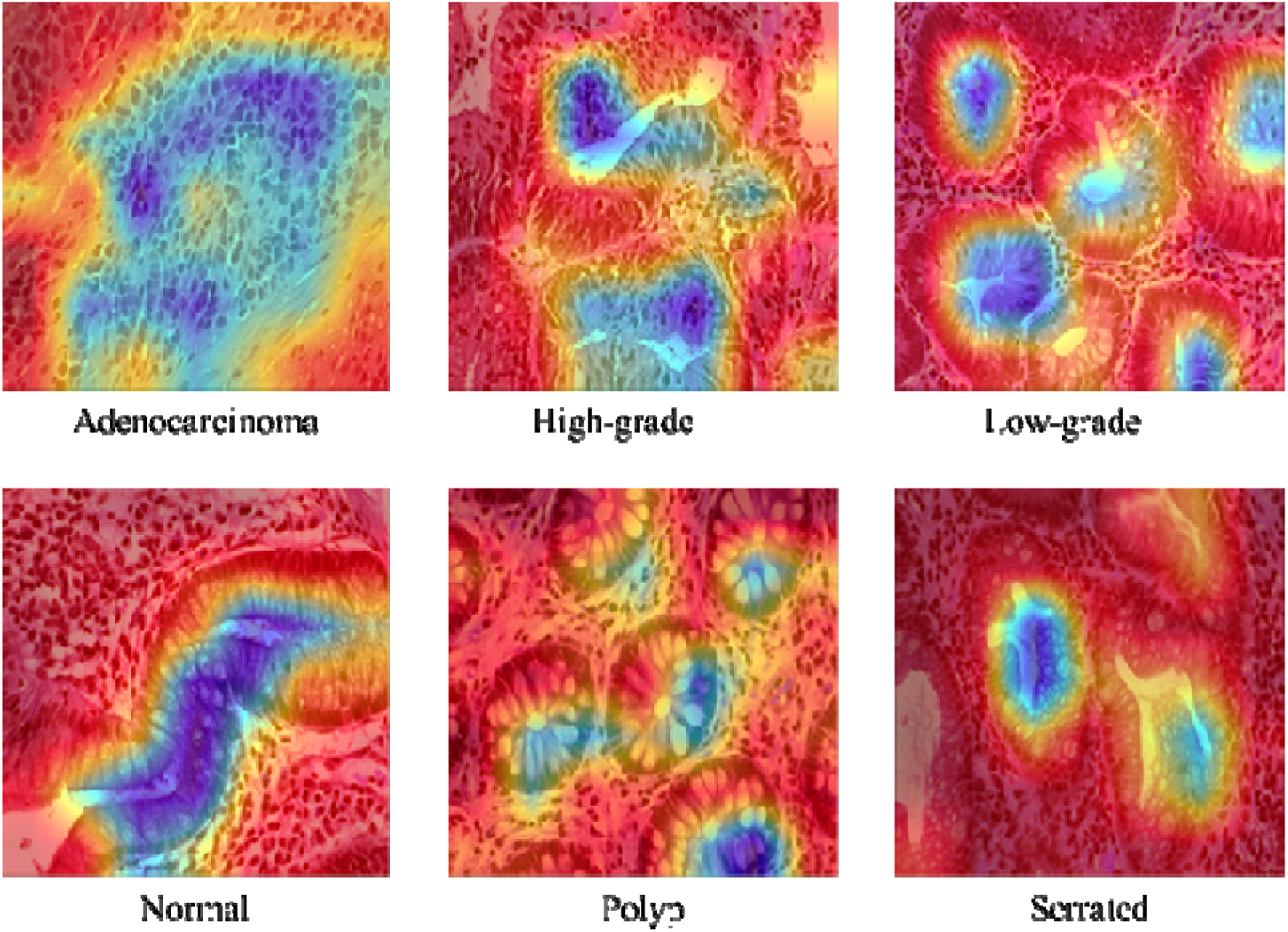
ResNet model GradCAM explanation on EBHI-Seg dataset.

**S. FIGURE 8.**
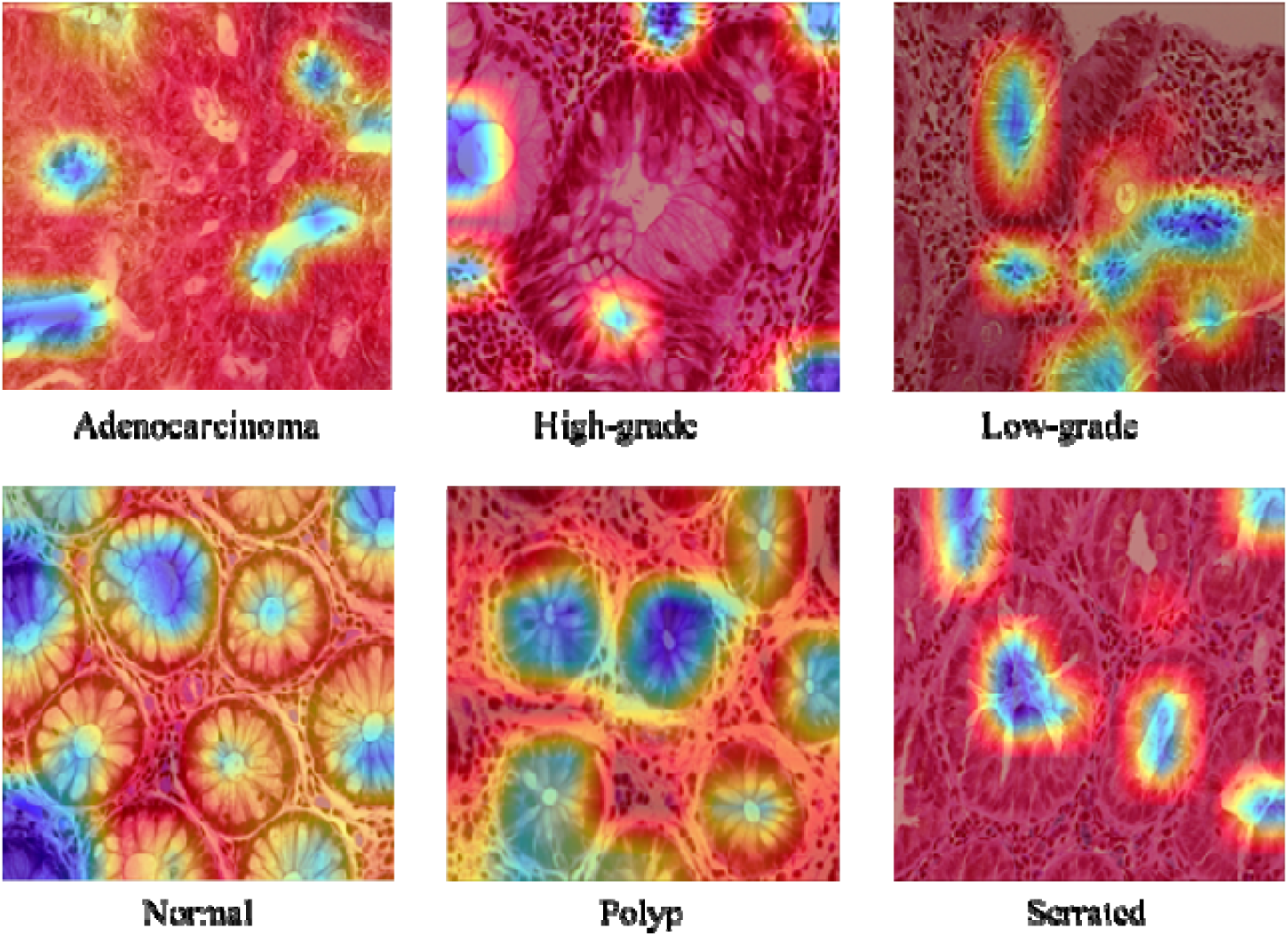
Swin Transformer model GradCAM explanation on EBHI-Seg dataset.

**S. TABLE 1.** Comparison of models in size (Params) and FLOPs.

| Model | Params (Million) | FLOPs (GFLOPs) |
| --- | --- | --- |
| **ResNet 18** | 11.18 | 1.82 |
| **ResNet 34** | 21.29 | 3.68 |
| **ResNet 50** | 23.52 | 4.13 |
| **SwinV2-t-W8** | 27.58 | 4.37 |
| **SwinV2-s-W8** | 48.96 | 8.48 |

